# Detection of alpha synuclein seeding activity in tear fluid in patients with Parkinson’s disease

**DOI:** 10.1101/2025.01.16.633314

**Authors:** Sezgi Canaslan, Matthias Schmitz, Fabian Maass, Peter Hermann, Susana da Silva Correia, Shuyu Zhang, Christoph van Riesen, Piero Parchi, Paul Lingor, Inga Zerr

## Abstract

Detection of alpha-synuclein seeding activity in tear fluid (TF) might provide a promising non-invasive biomarker for Parkinson’s disease (PD) diagnosis.

In this study, we applied the alpha-synuclein seeding amplification assay (aSynSAA) to detect misfolded alpha-synuclein (aSyn) aggregation in TF from PD patients. The discovery cohort included 11 PD patients and 13 controls, and the validation cohort consisted of 9 PD patients and 11 controls without synucleinopathies.

The aSynSAA yielded positive results in over 55% of PD patients. These findings were confirmed in a second cohort, including patients with prion diseases as a negative control for synuclein pathology. Our results demonstrate for the first time the ability of aSynSAA to distinguish between PD and control groups in TF, with PD showing the highest seeding activity compared to prion disease and control groups. Further comparisons between cerebrospinal fluid (CSF) and TF samples from the same individuals revealed consistent seeding results across both biofluids. These findings highlight the potential of tear fluid as a novel, accessible medium for detecting Lewy body-specific misfolded synuclein aggregation in PD, which could aid in early diagnosis and disease progression monitoring.

## Introduction

The novel paradigm-shifting technology of protein amplification assays is based on induced protein misfolding and is extremely sensitive ^1^. The amplification assays for synuclein (aSynSAA) have been successfully applied to brain and skin tissue and CSF samples in synucleinopathies ^2,3^. The method can potentially be extended to other accessible matrices, such as saliva and blood and was used to identify the underlying pathology even preclinically ^4,5^. Olfactory mucosa and saliva have already tested positive by aSynSAA ^6,7^. There is also data available on exosomes from blood containing misfolded proteins ^8,9^. However, complex matrices such as blood (and potentially oral and nasal swabs, too) bear the intrinsic problem of containing reaction inhibitors and other technological problems as it has been demonstrated for CSF ^10^. Tear fluid (TF) is a largely cell- and contamination-free bio fluid, which can be accessed non-invasively and with little discomfort for the patient. In addition, innervation of the lacrimal gland originates in the brainstem – a structure with is affected early in PD. We therefore selected TF as a testing platform for our experiments.

aSynSAA has been applied to detect misfolded prion protein in tear fluid in prion diseases being positive before the onset of clinical disease with similar diagnostic accuracy as for CSF^11^. α-Syn is detectable in tear fluid of synucleinopathies, with increased abundance in PD ^12^. In addition, there is evidence that α-Syn oligomers are detectable in saliva by asynSAA ^7^. We applied this powerful technology to the asynSAA to amplify α-Syn seeds in tear fluid from patients with PD and controls.

## Material and Methods

### Patients

Tear fluid samples were obtained from the Movement Disorder Biobank at the Department of Neurology, University Hospital, Göttingen, Germany. Patient samples were collected to facilitate prospective research projects related to Movement disorders. Parkinson’s disease patients were diagnosed according to the Movement Disorder Society (MDS) criteria ^13^. Control subjects with comparable age and sex, without clinical signs of neurodegeneration, were also selected from the biobank and consisted mostly of patients with peripheral neurological disorders like polyneuropathy or muscular disorders plus one patient with subcortical arteriosclerotic encephalopathy. The patients underwent neurological examination and history was taken by movement disorder specialists. Patients were eligible regardless of disease duration or severity. Ophthalmological comorbidities and topical ocular medications were recorded. Only those with topic hyaloron administration were allowed. Approval from local ethics committees was obtained (Ethics Committee of the University Medical Centre Göttingen, no. 37/11/21). Control samples from patients with prion disease were selected from the biobank of the German National Reference Center for Transmissible Spongiform Encephalopathies and donated tear fluid as part of a surveillance and biomarker discovery study (Ethics Committee of the University Medical Centre Göttingen, no. 11/11/93). The diagnoses were based on clinical criteria and *PRNP* genetic testing. Cerebrospinal fluid samples were obtained from the Movement Biobank, the prospective clinical study on cognitive decline in Parkinson’s disease (PARKA) and the Nerochemistry lab of the Department of Neurology. All studies comply with the Code of Ethics of the World Medical Association (Declaration of Helsinki).

### TF collection

Tear collection was performed as previously described ^12^. Briefly, tear samples were collected from both eyes using Schirmer test strips (Optitech, Allahabad, India). The strips were placed on the lower lid margin for 8 minutes and the wetting length was noted for each side. Patients with corneal inflammation or corneal ulcers were excluded. No topical anaesthesia was used for sample collection. Samples were frozen immediately after collection and stored in polypropylene tubes at -80°C until further analysis.

### Extraction of tear fluid

For the extraction of TF from the strips, TF extraction buffer was prepared in a final concentration of 40 mM PB pH 8.0, 170 mM NaCl and 0.0015% SDS. According to the wetting length of the strip, the required buffer volume was calculated as previously reported ^11^ (15 mm requires 50 microliters TF extraction buffer). If the wetting length exceeded 15-20 mm, the strip was cut into 2-3 pieces depending on the total length using sterilized scissors, which were cleaned with 70% ethanol between each use to prevent contamination. After incubation on the shaker at room temperature for one hour, strips were transferred to 0.5 ml Eppendorf tubes, from which the bottom part was cut. These tubes were inserted into 1.5 or 2 ml Eppendorf tubes to collect the tear fluid. Samples were centrifuged at 13.000 rpm for 20 minutes. The extracted tear fluid was collected and stored at -80 °C.

### SynSAA protocol

The aSynSAA reactions are conducted following the established protocol outlined by Groveman et al with minor changes ^14^. The His-tagged α-Syn protein was purified using His-affinity chromatography and later on, anion exchange chromatography as stated in Groveman et al. ‘s work with some additional changes ^14^. The assay was performed on black 96-well plates with clear bottoms. Initially, each well was loaded with 6 × 0.8 mm molecular biology grade silica beads using a bead dispenser. The lyophilized recombinant α-Syn is resuspended into 520 ml of 40 mM phosphate buffer (PB) pH 8.0 and resuspension is filtered using a 100 kDa Amicon filter for 6 min at 10.000 rpm to remove potential oligomeric α-Syn. For the reaction mix, 85 μl were prepared per well, containing a final concentration of 40 mM PB pH 8.0, 170 mM NaCl, 10 μM Thioflavin-T, 0.0015% SDS, and 0.1 mg/ml of recombinant α-Syn in 100 μl. After adding 85 μl of the reaction mix to each well 15 μl of extracted TF sample or CSF were pipetted into the wells. Samples are tested in triplicates except for 6 samples (Table 2,*). The plate was sealed with a transparent plate sealer and placed in a Fluostar Omega plate reader. The incubation was carried out at 42 °C with double orbital shaking at 400 rpm for one minute, followed by a one-minute rest period. Fluorescent measurements were taken every 45 minutes using 450 nm excitation and 480 emission filters for up to 30 or 110 hours for CSF and TF, respectively. The sample was considered positive when the fluorescence intensity reached at least two-fold of its initial value, measured 45 minutes after the start of the assay for TF and at least three-fold for CSF.

**Table 1.** Demographic and clinical characteristics of the cohorts.

|  | Age | Gender(F,M) | Modified H&Y stage | UPDRS III | Disease duration |
| --- | --- | --- | --- | --- | --- |
| PD | 71.2±8.7 | 4;16 | 3.1±0.9 | 34.8±12.3 | 8.7±6.6 |
| Controls | 69.0±9.7 | 5;13 | NA | NA | NA |

**Table 2.** Information about the SAA results on analyzed cohorts

|  | Negatives<br>(0 or 1 out of 3) | Positives<br>(2 or 3 out of 3) | Total<br>(n) | Percentage of Positivity (%) |
| --- | --- | --- | --- | --- |
| *Cohort 1* |  |  |  |  |
| PD | 5 | 6 | 11 | 54.54 |
| Controls | 12 | 1 | 13 | 8.00 |
| *Cohort 2* |  |  |  |  |
| PD | 4 | 5 | 9 | 55.55 |
| Controls | 5 | 0 | 5 | 0 |
| Prion Disease | 6 | 0 | 6 | 0 |
| *All* |  |  |  |  |
| PD | 9 | 11 | 20 | 55.00 |
| Controls | 17 | 1 | 18 | 5.56 |
| Prion Disease | 6 | 0 | 6 | 0 |

For the synSAA TF assay, we only used strips with a wetting length of more than 10 mm (preferably 15 mm), corresponding to approximately 10 μL of TF. Since shorter wetting lengths gave unequivocal results, they were then excluded from the analysis.

## Results

To develop a non-invasive diagnostic test for PD, we subjected TF from PD patients and controls to synuclein seeding amplification assay (asynSAA) analysis over a 110-hour period. PD samples showed a marked signal response (Fig. 1A). In contrast, control samples from patients with prion diseases and non-neurological controls exhibited no seeding activity except one control (Fig. 1A, Tab 1)). This control was diagnosed with advanced subcortical arteriosclerotic encephalopathy (SAE) with mild cognitive impairment (MCI), but no parkinsonism. Consequently, the presence of a subclinical aSyn pathology could not be fully excluded.

**Figure 1.**
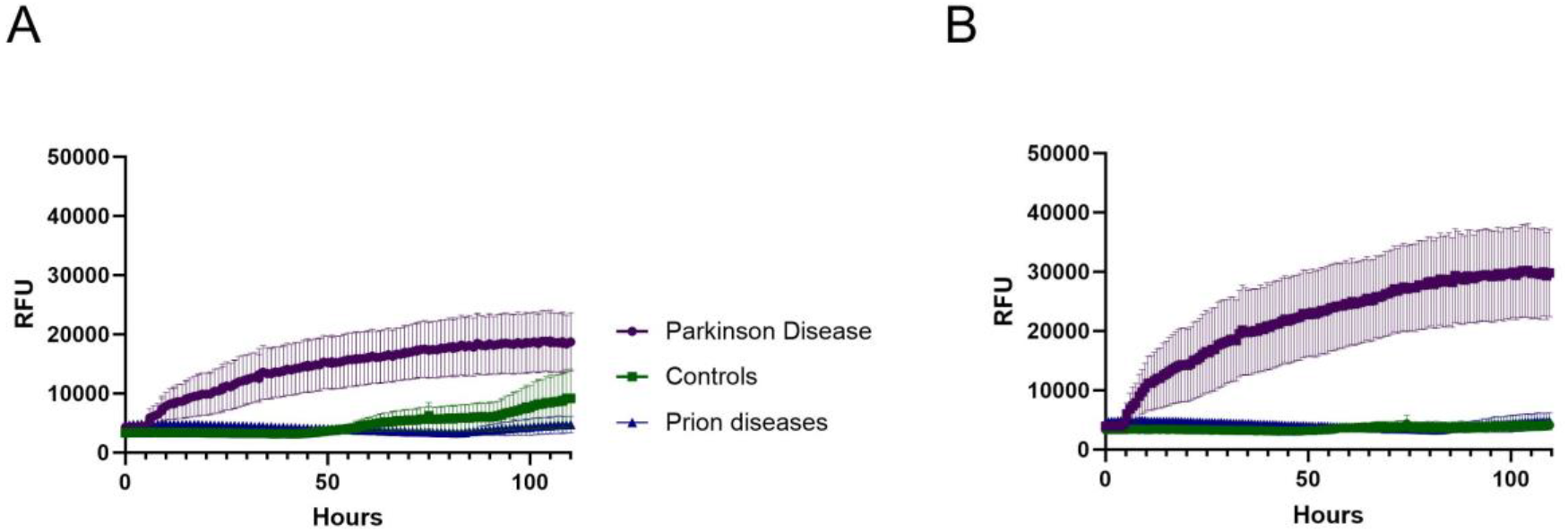
Comparison of seeding performance in TF among Parkinson’s Disease, Prion diseases, and control groups using SAA. A) The data represents the mean seeding activity indicated in relative fluorescence units (RFU) for each sample, with the solid line indicating the overall mean of the sample means for each group. Error bars indicate the standard error of the mean (SEM). (Parkinson’s Disease n=20, Controls n=18, Prion Disease n=6) B) The mean seeding activity for each group is represented in relative fluorescence units (RFU). The solid lines show the overall mean of true positive sample means (PD group-purple line), true negative samples (controls-green line) and prion disease samples (blue line). Error bars lines indicate SEM.

The diagnostic accuracy in our discovery cohort, consisting of 11 PD patients and 13 controls, showed that asynSAA was positive in 55% of PD samples, when defined by a signal increase of more than 50% and at least two-thirds of reactions positive (Table 2).

To gain a better understanding, we separately analyzed PD samples with positive results and control samples with negative results as shown in Figure 1B. The control curve remained flat, as expected, while the Parkinson’s disease curve became steeper and reached a higher fluorescence value.

To validate these results, we analyzed a second independent cohort, including samples from patients with prion diseases unrelated to α-Syn pathology that served as an additional control. Our findings were confirmed in the second cohort (Table 2).

### Comparison of seeding activity in CSF and TF

We then aimed to compare the seeding activity of α-Syn in SAA reactions seeded with cerebrospinal fluid (CSF) and tear fluid (TF) from the PD cohort. Reactions were seeded with 15 μL of CSF or 15 μL of TF obtained from the PD patients and the data are displayed in Figure 2. Although TF-seeded reactions were monitored over 110 hours, data up to 30 hours were analyzed to allow better comparison with CSF. The kinetic curves differed significantly, with CSF samples displaying a steeper curve compared to TF samples. Additionally, the average maximum fluorescence value of CSF was approximately four times higher than that of TF for the first 30 hours. Due to the limited number of matching CSF and TF samples, we included additional samples to obtain a more comprehensive understanding of the seeding kinetics of α-Syn. Figure 2B represents the true-positives samples from Parkinson’s disease cohort in CSF and TF.

**Figure 2.**
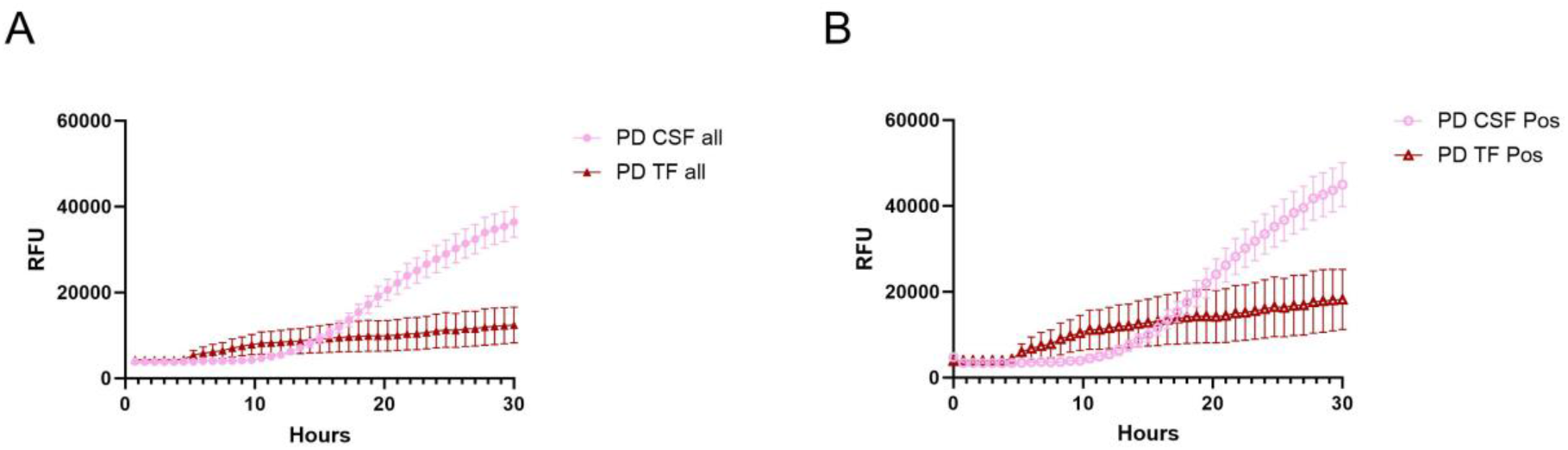
Comparison of CSF and TF seeding performance of individuals with Parkinson’s disease. The data represents the mean seeding performance for each sample, with the solid line indicating the overall mean of the sample means for each fluid. Error bars indicate the standard error of the mean (SEM). (CSF and TF Parkinson’s disease n=163 and n=20 for Figure 2A; CSF and TF Parkinson’s Disease true positive samples n=101, n=11 for Figure 2B)

## Discussion

For decades, research on biomarkers for neurodegenerative diseases was classically focused on CSF as a matrix because of its close relationship to the brain. In recent years, significant progress has been made concerning the development of blood-based assays, specifically in the field of Alzheimer’s disease and motor neuron diseases (Abeta, Tau, p-Tau, NFL) ^15,16^. The RT-QuIC/SAA technology goes even beyond and allows the detection of misfolded proteins in saliva and serum (in PD) and even in tear fluid (in prion disease) ^7,17,18^. The advantage of the SAA is the extreme stability of misfolded protein aggregates at various conditions compared to protein-based assays. The technology has already entered the field of clinical diagnosis in prion diseases and has been highlighted as one of the best available CSF biomarkers for synucleinopathies ^19^.

For synucleinopathies, synSAA has been proposed as a potential biomarker ^20, 21^. SynSAA has already proven effective for qualitative α-Syn detection in the CSF at the preclinical and early clinical disease stage ^3-5,22,23^ and biological definitions and staging systems of PD based on the presence of α-Syn pathology have been suggested ^20,21^.

The use of tear fluid as a matrix for biomarker analysis has multiple advantages over other body fluids. The collection is non-invasive and can be performed repeatedly, even by trained non-medical personnel. It is safe and can be applied in a non-hospital setting and it might therefore be very suitable for screening and monitoring purposes. It is free of contamination by blood and blood particles, as frequently occurring in CSF samples and may obscure the results. The composition of TF might be even more suitable than CSF, where the intrinsic inhibitors might affect the test results ^10^.

Testing tear fluid offers a unique opportunity for biomarker development, specifically for emerging technologies that are based on aggregation assays. Unlike surrogate biomarkers commonly proposed for many neurodegenerative disorders, which lack specificity, aggregation assays (SAA) are based on the structural characteristics of a misfolded protein. These altered aggregation characteristics of the disease-related misfolded protein are used to amplify the seeds of the misfolded proteins of interest in a prion-like manner in vitro, followed by detection of the newly formed aggregated species (Real-Time Quaking-induced Conversion, RT-QuIC, asynSAA). This approach identifies tiny amounts of the misfolded species at the femtomolar level and is ideally suited to detect synuclein aggregation activity (synSAA) in tear fluids.

In our study, we demonstrated aSyn seeding activity in TF derived from PD patients via asynSAA. Using two independent cohorts—a discovery cohort and a validation cohort—we observed an overall diagnostic accuracy of 55% for PD diagnosis. Of the 24 controls, only one TF sample was diagnosed as MCI due to SAE revealed a positive signal response in the SAA, which is in line with current evidence from the literature, where underlying preclinical synuclein pathology is suggested in various clinical scenarios. Of interest, the aggregated kinetic curves from TF and CSF show slightly different dynamics with a steady increase for CSF and flattening of the curve in TF. One potential explanation might be the different level of synucelin aggregates as a substrate for seeding, another might be diverse synuclein species in both biofluids, which then react in a different way. We also observed that some PD patients had a very fast response in TF, even before CSF. Since the numbers are low, we cannot draw definite conclusions, but we speculate that disease stage or disease subtype might play a role.

However, our study has several limitations. Samples were derived from one center only and therefore confirmatory analyses including samples from multiple centers are necessary. In CSF, a pooled sensitivity and specificity to differentiate synucleinopathies from controls with αSyn-SAA was indicated with 0.88 (95% CI, 0.82–0.93) and 0.95 (95% CI, 0.92–0.97) ^2^. Our data shows a diagnostic accuracy of 55%, which may be due to our current asynSAA protocol and further protocol modifications may be necessary to achieve similar results in the TF asynSAA. However, this result may also indicate that only a subgroup of PD patients shows aSyn seeding activity in their TF. Correlative analyses with corresponding CSF samples in larger cohorts, including subgroups of PD patients with different clinical characteristics (early vs. late onset, tremor vs. bradykinetic onset, PD with and without RBD, etc.) will be informative here.

In neurodegenerative diseases, the accumulation of protein aggregates in the brain starts years before the onset of symptoms. Timely identifying α-Syn accumulation is crucial for diagnostic purposes and early evaluation of clinical trajectories. Moreover, individuals with pre- or early clinical disease are putative candidates for trials assessing the efficacy of disease-modifying drugs and the main target of future therapeutic interventions. The development of an easily applicable assay in tear fluids would be an important milestone towards this goal.

## Data availability

All data are available upon reasonable request

## CONTRIBUTORS

MS, PL and IZ conceived the study, SC performed experiments, SC, MS, IZ analysed data, interpreted results and performed statistical analyses. MS, FM, PP and IZ acquired data and contributed to technical expertise. FM, CvR, SC,PH, SZ, PL acquired biological samples. SC, MS, IZ drafted the manuscript. All authors read and approved the manuscript.

## Compliance with ethical standards

Conflict of interest: The authors declare that they have no conflict of interest.

